# MiRNA and Erythropoietin profiles during the menstrual cycle in relation to hematological and lipid biomarkers

**DOI:** 10.1101/2021.11.05.467468

**Authors:** Helena Bergström, Carmel Heiland, Linda Björkhem-Bergman, Lena Ekström

## Abstract

**Background:** Circulatory micro RNAs (miRNA) have been discussed as complementary diagnostic markers in cardiovascular diseases, and in anti-doping testing. MiR-144 and miR-486 have been associated with cholesterol homeostasis and hematopoiesis, respectively. In addition, they have been suggested as putative biomarkers for autologous blood transfusion and erythropoietin (EPO) doping. The aim of the present study was to assess the variability of miR-144-3p/5p, miR-486-5p/3p and EPO during the menstrual cycle. Secondary aim was to study the correlations between miRNAs, EPO and hematological parameters and lipids.

**Methods:** 13 healthy women with regular menses were followed with weekly blood sampling during two whole menstrual cycles. MiRNAs were analyzed using TaqMan and PCR followed by calculation of the relative expression for each miRNA using ddCT approach.

**Results:** There was no menstrual cycle variability in miRNAs and EPO. MiRNA-144-3p was associated with HDL-C (rs=-0.34, p=0.036) and miRNA-486-5p with Hb (r_s_=0.32, p=0.046). EPO concentrations correlated to lymphocytes (r_s_=-0.062, p=0.0002)_),_ Hb (r_s_= -0.42, p=0.0091), HDL-C (r_s_=0.36, p=0.030) and triglycerides (r_s_=-0.54, p=0.0006).

**Conclusions:** The results of this study may increase the understanding of how miR486-5p and miR144-3p as well as EPO correlate to hematopoietic and lipid biomarkers.

## 1. Introduction

Micro RNAs (miRNAs) are short (−22 nucleotides) non-coding RNAs that post-transcriptionally regulate gene expression through binding and silencing of complementary messenger RNAs (mRNA). miRNAs are relatively stable and can be reliably quantified in extracellular body fluids e.g. serum or plasma [1]. The miRNAs may be present in two isoforms, the 3′ and the 5′ strand, and usually one of the strands is the referentially active [2]. In the circulation, the miRNAs are protected from ribonucleases by being encapsuled in exosomes [3], micro-vesicles [4] or lipoproteins [5]. The levels of miRNAs may be dependent on ethnicity and age [6, 7]. Also, as gender differences exist, it has been suggested that some miRNAs are hormonally regulated [8, 9].

Different miRNAs have been identified as diagnostic markers for cardiovascular diseases/disorders such as acute myocardial infarction (AMI) [10], atherosclerosis [11] [12], and dyslipidemia [13]. MiRNA-144 is found in the circulation and it has been associated with cholesterol homeostasis [12]. Studies suggest that miRNA-144 regulate the cholesterol metabolism via adenosine triphosphate-binding cassette transporter A1 (ABCA1) suppression [12, 14]. Higher plasma levels of miR-144-3p have been associated with the presence, as well as severity of cardiovascular disease (CVD), and proposed as predictors of AMI [15] [16]. Another putative cardiovascular miRNA biomarker includes miR486-5p, where circulatory levels have been associated with hematopoiesis [17] and AMI [18]. In contrast to miR144-3p, that is predominantly found in plasma, miR486-5 is also highly expressed in red blood cells [1, 19].

MiR-144-5p has been reported to be up-regulated after erythropoietin EPO administration [20] whereas miR-144-3p, miR-486-5p and miR-486-3p increased after autologous blood transfusion (ABT) [21]. To increase the sensitivity and detection window for blood doping i.e. misuse of erythropoietin stimulating agents (ESA) and blood transfusions, it has been suggested that miRNAs could be included as complementary biomarkers in Athlete Biological Passport (ABP) [22]. However, the intra-and interindividual variability of miR-144 and miR-486, has not been investigated in fertile women.

Recently, it has been suggested that mutations in hematopoietic stem cells in the bone marrow, a condition known as clonal hematopoiesis of indeterminate potential (CHIP), contribute to the process of atherosclerosis and CVD [23]. Also, an association between EPO and the size of aortic abdominal aneurysms, was found in humans [24]. Interestingly, the expression of the EPO receptor was upregulated when another miRNA associated to CVD, miR-135b was downregulated. This was found to increase the stability of the atherosclerotic plaque in mice [25]. Subsequently, the link between EPO and biomarkers of atherosclerosis warrants further investigation.

To our knowledge, there are no studies where miR-486, miR-144 and EPO have been measured longitudinally in healthy, regularly menstruating subjects. This prompted us to perform a pilot study in female volunteers with the primary aim to assess the variability of miR-144-3p/5p, miR-486-5p/3p and EPO, during the menstrual cycle phases. The secondary aim was to study the correlations between circulatory levels of miRNAs, EPO and hematological parameters and lipids.

## 2. Material and methods

### 2.1 Study population

This is a sub-study of a single-center, observational study, including 17 healthy women with regular menstrual cycles that were monitored for two cycles in a row, with blood samples taken once a week. The study design and demographics of the participants have been previously described [26]. The variability of hematological and lipid biomarkers during the menstrual cycle has also been described in our cohort [27],[26], [28].Serum was immediately prepared (spinning 3500 g for 15 min) and stored at --80 C prior to microRNA and EPO analyses. To identify the exact menstrual phase for each sample, hormone concentrations of estradiol, progesterone, luteinizing hormone (LH) and follicular stimulating hormone (FSH) were analyzed at Division of Clinical Chemistry using immunological methods as previously described [28].

For the present study serum samples from 15 women were available, contributed with all together 34 samples. For analyses of EPO, six subjects had samples available from all three menstrual phases (n=18), and seven subjects had samples from two phases (n=14) (three with luteal and ovulation, two luteal and follicle and two with ovulation and follicle). Two subjects contributed with only one sample each (one follicular and one luteal). The two subjects with only one serum sample available were excluded, and two samples failed the micro-RNA preparation (absence of PCR signals of control miRNAs) and thus, the total number of samples for miRNA analyses was 31.

### 2.2 Micro RNA analyses

Micro RNA was extracted from 200 uL serum samples using miRNeasy Serum/Plasma Advanced Kit (Qiagen, Hilden, Germany) according to the manufacturer’s manual. No visual signs of hemolysis could be seen in the serum samples. The micro-RNA samples were subjected to cDNA conversion using Thermo Fisher cDNA TaqMan™ Advanced miRNA cDNA Synthesis Kit and the cDNA were diluted ten times according to protocol. The diluted cDNAs were used as templates in premade TaqMan assays targeting the miRNAs of interest, i.e., miR-144 (IDs 477913_mir/477914_mir, Life Technology) and miR-486 (assay ID 478128_mir/478422_mir) using TaqMan™ Fast Advanced Master Mix. The real-time polymerase chain reaction (PCR) was conducted on StepOne and the relative expressions for each miRNA were calculated using the ddCT approach, with one sample as calibrator [29]. For comparison of expression levels between the different miRNAs, one sample of miR-486-5 was used as calibrator. MiR26b-5p (assay ID 478418-mir) and miR27b-3p (ID 478418-mir) were used as control “house-keeping genes” as recommended by the provider. Finally, the relative expressions of the miRNAs were calculated using only miR-26b as a control as well as using the mean cycle threshold (Ct) from both miR26b and miR27b. Ct-values >35 were considered as undetectable.

### 2.3 EPO analyses

The EPO concentrations from 34 serum samples were determined using the Human Erythropoietin ELISA Kit from Stemcell Technologies. Briefly, 50 µL of serum, analyzed in duplicate, Buffer A, and biotinylated anti-EPO antibody, were added to microplate wells and incubated for 2 hours at room temperature. The wells were washed, Streptavidin horseradish peroxidase conjugate added to each well, and the plate incubated for 60 minutes on the laboratory bench. After washing the wells again, tetramethylbenzidine substrate solution was added to each well, incubated for 15 minutes and stop solution was added. The absorbance was measured at 450 nm using SpectraMax Plus 384 Microplate Reader (Molecular Devices, LLC, San Jose, CA).

### 2.4 Lipid and hematological parameters

Cholesterol, TG, and HDL-C were analyzed by enzymatic assay followed by photometry as previously described [26]. Low-density cholesterol (LDL-C)was calculated according to Friedewald, Levy and Fredrickson [30]. Sysmex XN-1000 was used for the analyses of red blood cell (RBC) count, reticulocytes % (RET %), lymphocytes and hemoglobin (Hb).

### 2.5 Statistical analyses

For miRNAs and EPO, D’Agostino & Pearson normality test was conducted to test for Gaussian distribution of data and arithmetic means. None of the variables had Gaussian distribution. The inter-individual differences were evaluated using Kruskal-Wallis test. For the intra-individual variability, coefficient of variation (CV) was calculated. Levels of EPO and the relative expression of miR144 and miR-486 during the three menstrual phases (luteal, ovulation, and follicle phase) were compared using ANOVA test followed by Kruskal-Wallis multiple comparison test. Correlations between miRNAs/EPO and hematological and lipid parameters were evaluated using Spearman rank test, A p-value p≤0.05 was considered statistically significant. Analyses were performed in GraphPad Prism 8.3.0.

## 3 Results

### 3.1 Expression profile of miR144 and miR486

Both control miRNAs (miR26b and miR27b) were stable, and their Ct-values showed strong correlations (rs=0.78, p <0.0001), as the Ct-values did not differ between the menstrual cycle phases (data not shown) thus confirming the validity of the combined normalizing miRNA approach. There were no differences in the relative serum miRNA concentrations of the miR486-5p (p=0.91), miR144-3p (p=0.98), miR486-3p (p=0.34) and miR144-5p (p=0.26) between the three menstrual phases using Kruskal-Wallis test (**fig** 1).

**Figure 1:**
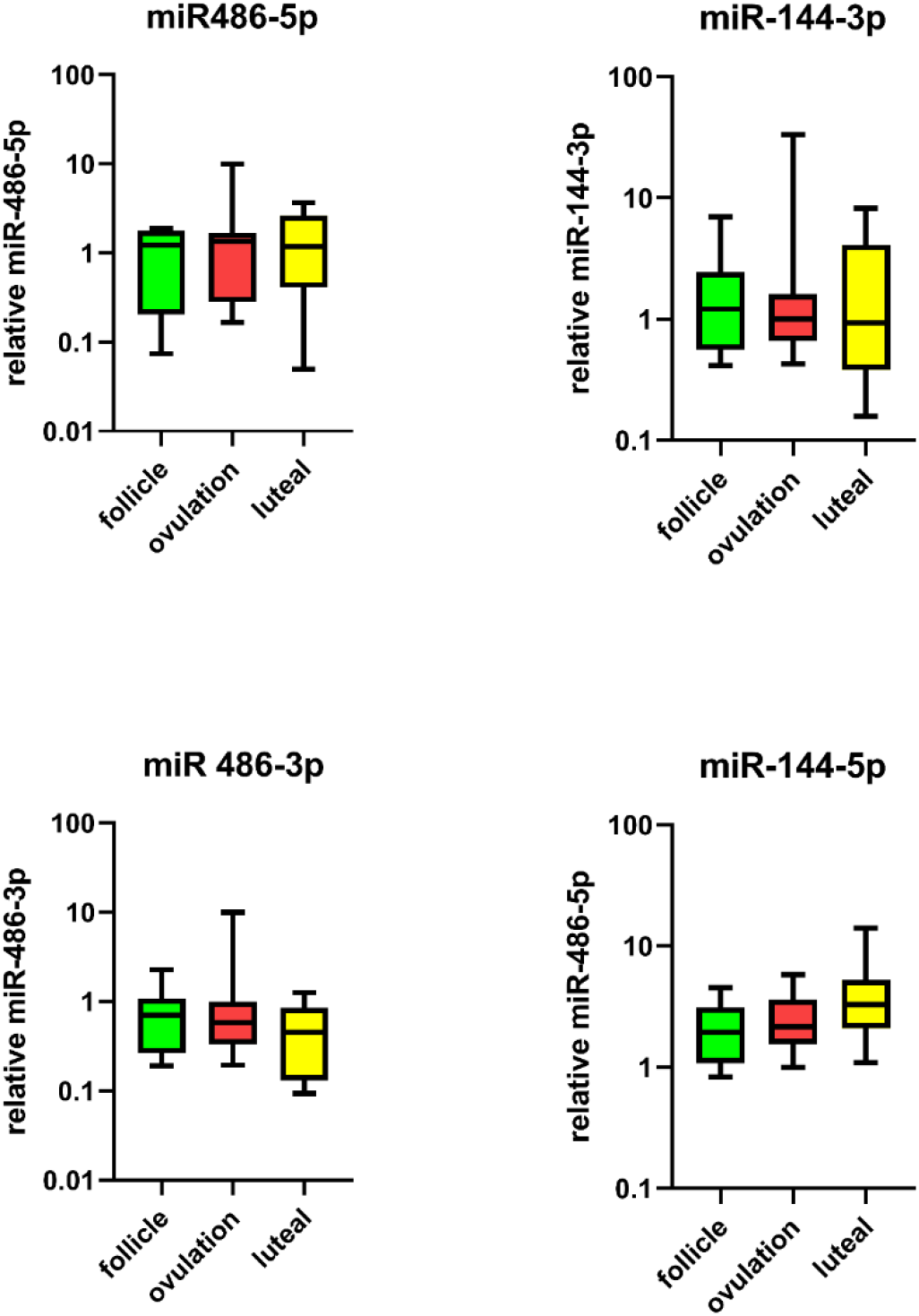
Relative levels of a) miR-486-5p b) miR144-3p c) miR486-3p d) miR-144-5p in the follicle (n=9), ovulation (n=11) and luteal phases (n=11) of the menstrual cycle in 13 healthy women.

In addition, the same results were found performing the statistical analyses using only miR26b as control, i.e., no differences between the menstrual phases were observed (data not shown).

There were strong correlations between miR-486-5p and miR-144-5p (r_s_=0.55. p=0.0005) and between miR-144-3p and miR486-3p (r_s_=0.62, p<0.0001). The correlation between miR486-5p and miR144-3p did not reach statistical significance (data not shown).

The relative expression of the different miRNAs shows, that miR486-5p and miR144-3p had a higher presence in serum as compared to the complementary miRNAs (**fig**. 2).

**Figure 2:**
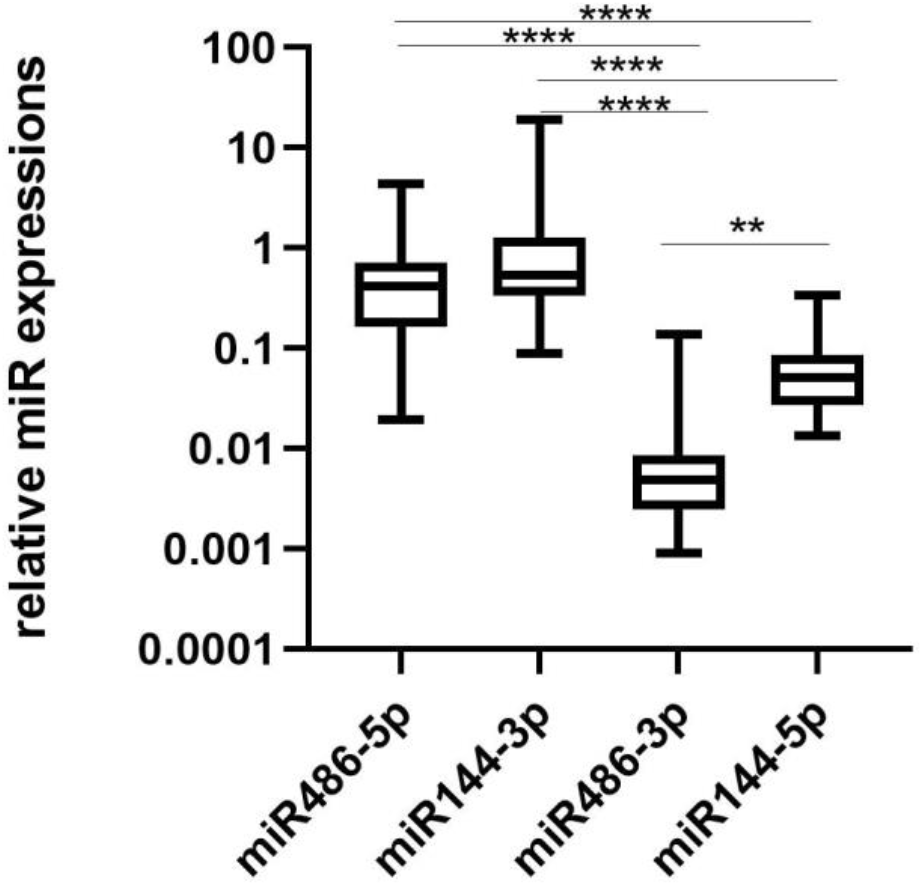
Relative expression of different miRNAs in the circulation in 13 women using a miR486-5p sample as calibrator. Number of samples include for miR486-5p (n=41), miR144-3p (n=42), miR486-3p (n=40), miR144-5 (n=31). **p<0.01, ***p<0.001, ****p<0.0001

The relative mean expression of miRNA-486-5p (mean 0.42) and miRNA144-3p (mean 0.53) were 100-fold and 10-fold higher than miRNA-486-3p (mean 0.005) and miR144-5p (mean 0.05), respectively.

The age of the participants correlated negatively with their mean expression of miR486-5p (r_s_=-0.70, p=0.0091) (**fig**. 3), but not with the other miRNAs investigated (data not shown).

**Figure 3:**
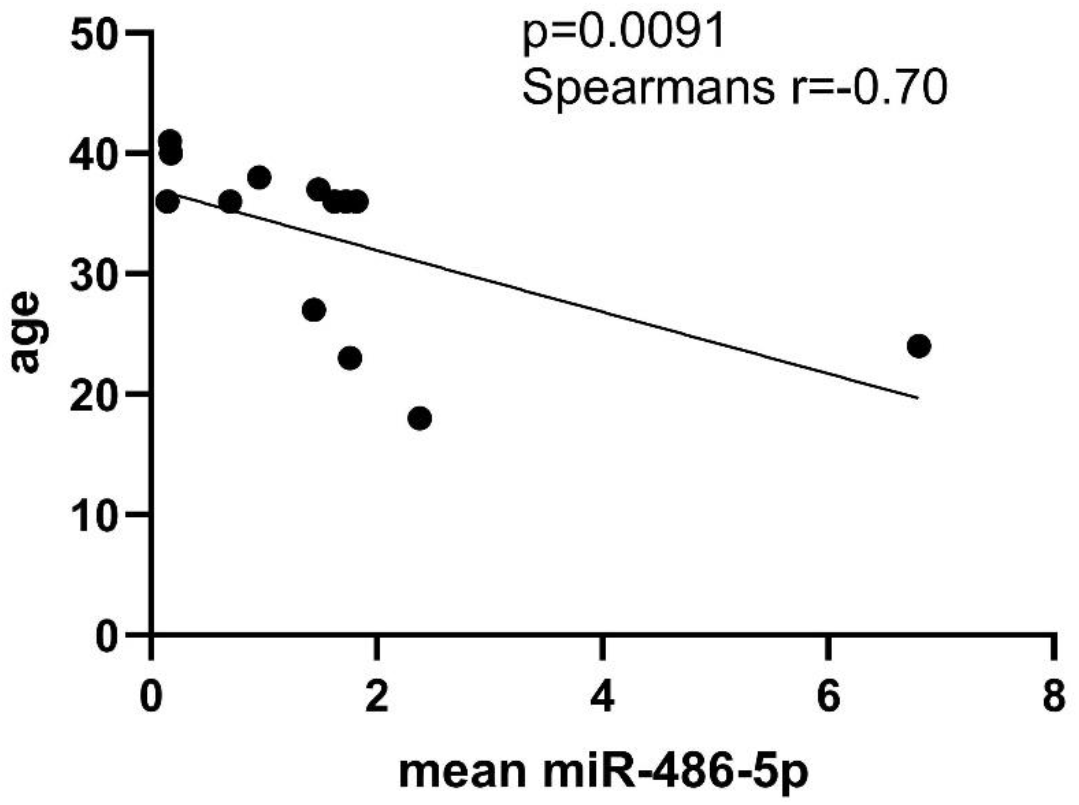
Correlation analysis between age and mean relative expression of miR-486-5p of the 13 participants.

There were large inter-individual variations in the abundance of the circulatory miR486-5p and miR144-3p, with 20-fold and 200-fold differences between the highest and lowest relative expression, respectively. The intra-individual median coefficient of variations (CV%) were 54.8 % (range 8-82 %) for miR486-5p and 45.7 % (range 3-120 %) for miR144-3p, respectively.

### 3.2 miR486-5p and miR144-3p correlation with hematological parameters and lipids

The miR486-5p correlated to Hb (r_s_=0.32, p=0.046) and reached borderline significance with RBC (r_s_=0.31, p=0.0549) and EPO (r_s_-0.31, p=0.0573) (**fig**.4).

**Figure 4:**
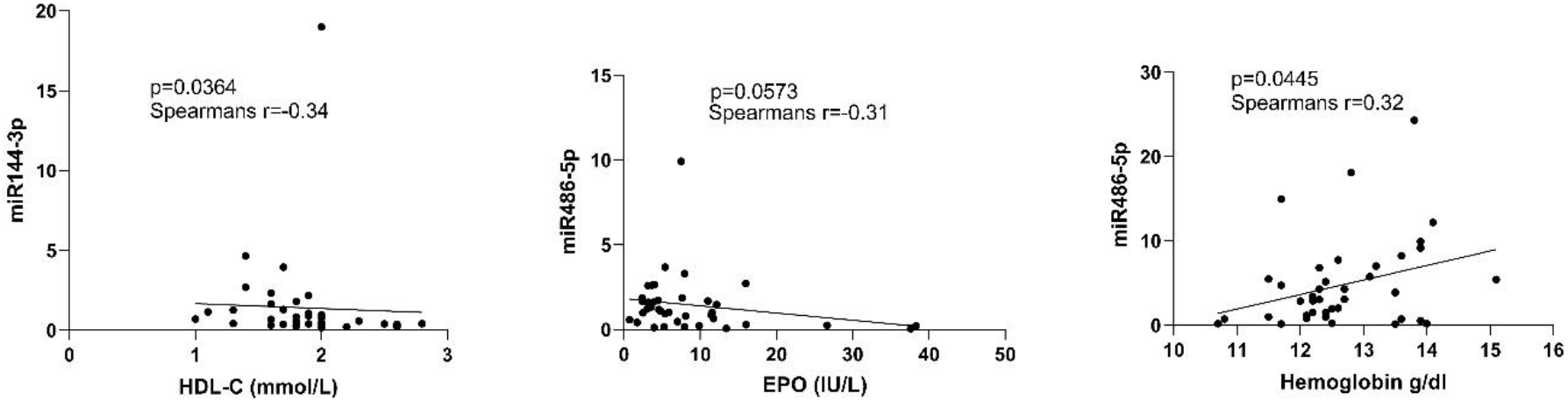
Correlation analyses between miR-144-3p and HLD-C, miR-486-5p and Erythropoietin (EPO) and hemoglobin (Hb) respectively.

There were no correlations of significance between the other hematological and lipid parameters, **table** 1.

**Table 1.** Correlations between hematological and lipid biomarkers and miRNAs levels in 31 samples collected through two menstrual cycles from 13 women.

|  | miRNA-486-3p | miR-486-5p | miR-144-3p | miR-144-5p |
| --- | --- | --- | --- | --- |
| RET%<sup>⌘</sup> | $r_s$=0.03 | $r_s$ = 0.21 | $r_s$=-0.05 | $r_s$=-0.0009 |
| RBC<sup>⌘</sup> | $r_s$=-0.007 | **$r_s$ = 0.31<sup>#</sup>** | $r_s$=-0.14 | $r_s$=0.06 |
| Hb<sup>⌘</sup> | $r_s$=0.14 | **$r_s$=0.32\*** | $r_s$=-0.09 | $r_s$=0.06 |
| EPO | $r_s$=0.2 | **$r_s$=-0.31<sup>#</sup>** | $r_s$=0.12 | $r_s$=0.08 |
| HDL-C<sup>⌘⌘</sup> | $r_s$=-0.06 | $r_s$=-0.07 | **$r_s$=-0.34\*** | $r_s$=0.03 |
| LDL-C<sup>⌘⌘</sup> | $r_s$=0.11 | $r_s$=0.12 | $r_s$=0.12 | $r_s$=0.10 |
| TG<sup>⌘⌘</sup> | $r_s$=0.22 | $r_s$=0.13 | $r_s$=-0.08 | $r_s$=-0.08 |

Spearman correlation coefficient ( $r_s$ ) presented. Significant data marked in bold text.
\* $p$ <0.05 # $p$ =0.05. ⌘ from Mullen et al ⌘⌘ in Bergström et al
RET%=reticulocytes, RBC=red blood cells, Hb=hemoglobin, EPO=erythropoietin, HDL-C=high density lipid cholesterol, LDL-C=low density lipid cholesterol, TG = triglycerides

Furthermore, the miRNA144-3p showed a significant negative correlation with HDL-C (r_s_-0.34, p=0.036) (**fig**.4) and no correlation with the other lipid biomarkers.

The correlation analyses were also conducted using only miR26b as a normalization control and the same results were found i.e., significant correlations between miR-486-5p and RBC, Hb and EPO and between miR144-3p and HDL-C (data not shown).

### 3.3 Serum EPO concentrations in relation to menstrual cycle phases, hematological parameters and lipids

There was no significant difference in the EPO concentration (n=34) between the three menstrual cycle phases (**fig**. 5).

**Figure 5:**
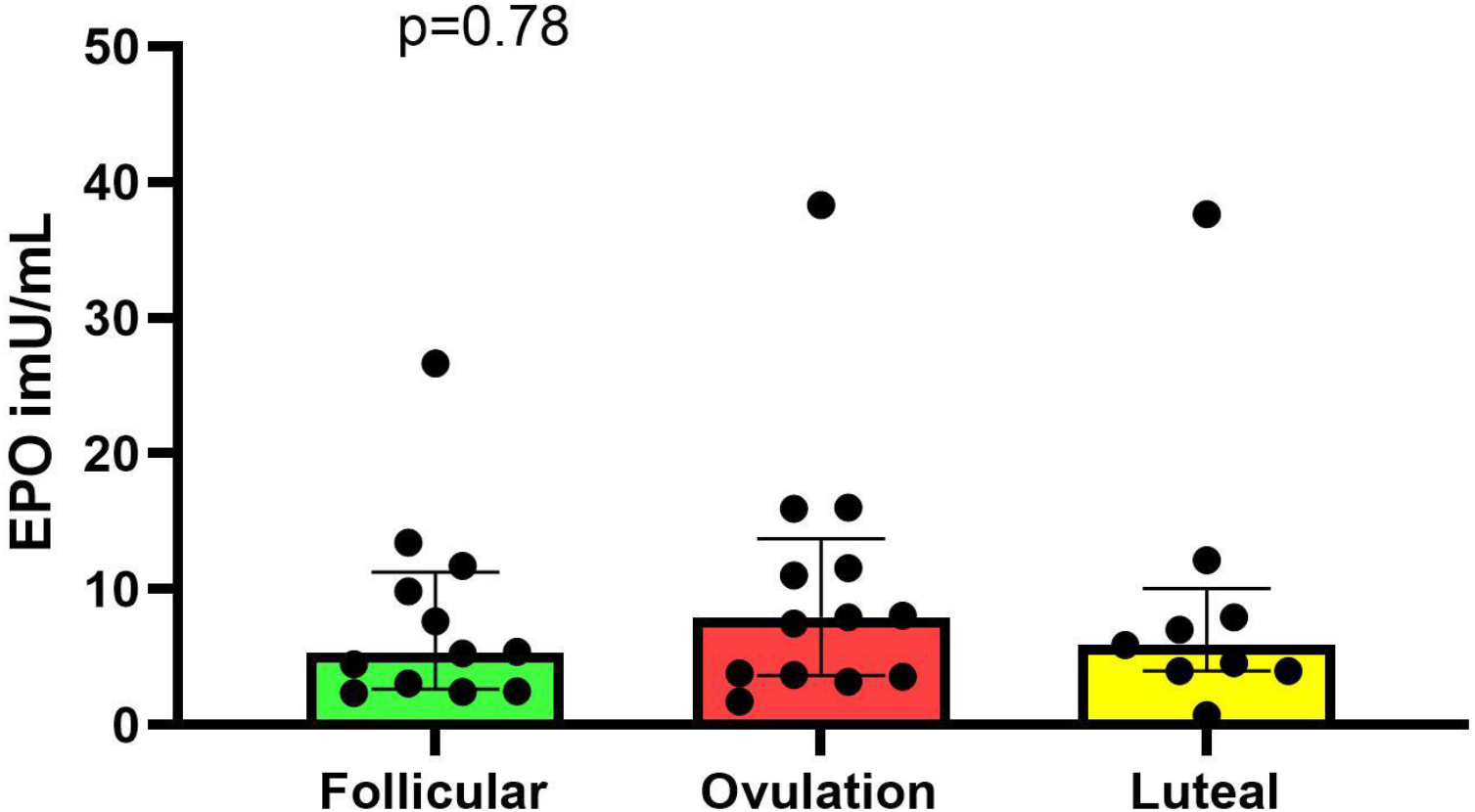
Variations in Erythropoietin (EPO) during the three menstrual phases of 15 healthy women. The total number of samples were 34 (follicle n=12, ovulation n=13 and luteal phases n=9)

There was significant inter-individual variability in EPO (p=0.0001) and the median (CV%) for intra-individual variability was 60 % (range 13-93 %). Furthermore, there was a significant correlation between EPO and lymphocytes (r_s_=-0.062, p=0.0002) (**fig**. 6 a). For EPO, negative correlations with Hb (r_s_=-0.42, p=0.009) (**fig**. 6 b) and TG (r_s_=-0.54, p=0.0006) (**fig**. 6 c) were found. Finally, a positive correlation was found between EPO and HDL (r_s=_ 0.36, p=0.030) (**fig**. 6 d).

**Figure 6:**
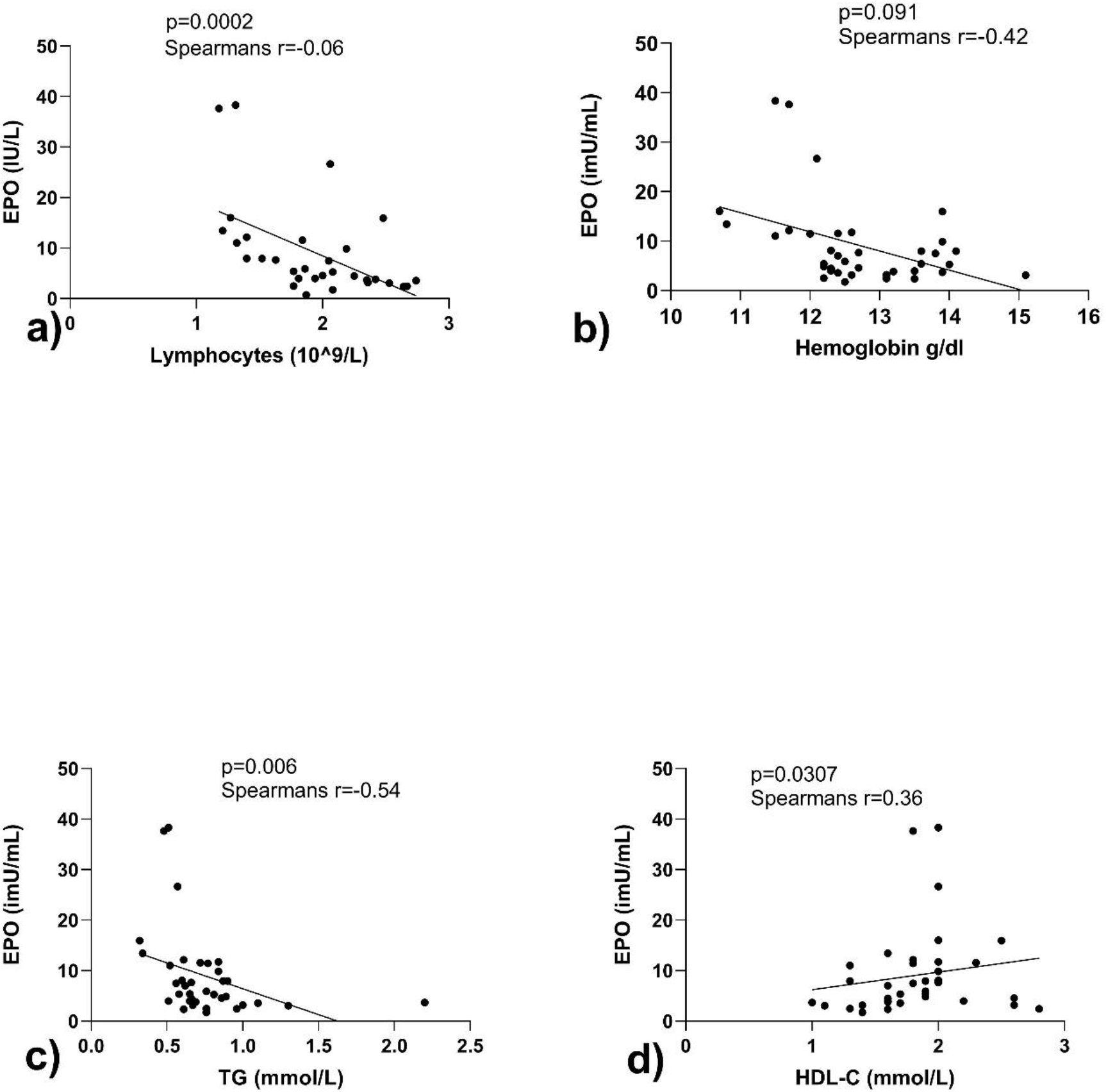
Correlations between Erythropoietin (EPO) and a) lymphocytes b) hemoglobin (hb) c) triglycerides (TG) and d) high-density cholesterol (HDL-C) during two menstrual cycles in 34 samples from 15 healthy women.

## 4. Discussion

### 4.1 Relative expression of miRNAs

The novelty of this study is, that for the first time miR144 and miR-486 have been longitudinally sampled, in relation to the menstrual cycle, on an individual basis. The lack of a menstrual cycle variability, is in agreement with a pooled analysis of circulatory miRNAs in nine women, including miR-144-3p and miR-486-5p [31]. However, there are studies, where the miRNA expression in endometrium have been associated with menstrual hormone fluctuation [8], but it is possible that tissue specific alterations in miRNAs are not reflected in serum [32].

The expression profiles of miR144-3p and 486-5p did not correlate in our participants, as opposed to a study by Wakabayashi et al, who discerned a weak correlation in serum from Japanese healthy men [6]. This may be due to gender and/or ethnicity differences between the study populations, as well as different methods applied. Both miR-144-3p and miR486-5p were found at considerably higher levels than their sequence complementary isoforms. In addition to being less abundant, these complementary miRNAs did not show any association to any of the biomarkers studied, implicating that their use as diagnostic biomarkers in CVD may be questionable.

### 4.2 miRNAs, lipids and hematopoiesis

In the present study, miR144-3p was negatively correlated with HDL-C, supporting the hypothesis that miR-144 is involved in the homeostasis of cholesterol. Some studies suggest that silencing of miR-144 could be a novel approach to targeting HDL levels by reversing cholesterol efflux [14]. Moreover, miR-144-3p has been suggested to regulate cholesterol homeostasis in colorectal cancer [12]. Although previous reports have indicated that miR144 is involved in erythropoiesis in humans [33], our results indicate no direct association with the hematological parameters investigated.

Furthermore, there was a correlation between miRNA-486-5p and red blood cells (RBC), Hb and EPO. This is expected, as miRNA-486-5p is highly abundant in erythrocytes [19, 34]. Even though the expression of miR-486-5p is high in erythrocytes, studies indicate that miR-486-5p is present in high levels in plasma samples [31], a prerequisite to function as circulatory miRNA biomarker. With this study, we confirm that miR486-5p indeed is present in serum, in similar range as miR144-3p. Additionally, it should be mentioned that HDL-C has been identified as a carrier of miR486-5p [35] However, there was no association between circulatory miR486-5p and HDL-C in our healthy participants. The reason for this could be that the HDL-C transport is elevated only in vulnerable coronary artery patients, as proposed [35].

### 4.3 EPO - variability, lipids and hematopoiesis

In our study, there was a large inter and intraindividual variability in EPO, and the levels of EPO correlated as expected with Hb [36]. However, we did not find any variability in EPO during the hormonal fluctuations throughout the menstrual cycle, confirming the results from a previous study in Japanese women [37].

Notably, EPO showed a positive correlation to HDL-C and a negative correlation to TG. This is in line with studies of treatment with recombinant human EPO (rhEPO), where decreases in LDL-C and TG, and increases in HDL-C have been shown [38, 39]. Furthermore, in patients with acute renal failure after coronary bypass surgery, the decrease in TG and total cholesterol negatively correlated with the increase in EPO [40]. It has been suggested that EPO activates lipid catabolism in peripheral adipose tissue, by activating the Janus Kinase 2 (JAK2)/STAT pathways, amongst others [41]. However, studies in vivo show that EPO does not activate lipolytic pathways in human white adipose tissue (WAT) [42]. Interestingly, studies in mice suggest that adipocytes in bone marrow are regulated by endogenous EPO, and that these adipocytes are of different origin and functionality than those found in WAT and brown adipose tissue (BAT) [43]. Furthermore, in obese mice with dyslipidemia, EPO administration induced distinct differential changes in bone marrow tissue, compared to normal mice, supporting a role for EPO in lipid metabolism [44].

In our study, there was a significant association between EPO and lymphocytes. Previous studies in mice have suggested that lymphoid cells lack the erythropoietin receptor [45], but recent evidence propose that EPO possesses important immune-modulating effects in humans [46]. Innate cells like monocytes can produce EPO and EPO has a modulating effect on T cells in the adaptive immune system [46]. At the same time, both B-cells and T-cell have been implicated in the process of atherosclerosis [47] Thus, it is possible that the association between EPO and other biomarkers of atherosclerosis may be indirect, with the common denominator being the immune system. Nonetheless, several studies have investigated the non-hematopoietic roles of EPO in normal physiology and metabolism [48].

### 4.4 miRNAs and EPO Doping

Clinical analysis of EPO is performed only in certain conditions but is of interest in anti-doping setting since EPO is a forbidden in sport. To detect EPO abuse, an electrophoretic method that separates endogenous EPO from recombinant EPO and erythropoietin stimulating agents is used, a method that lately has been improved in regards of detection window [49]. Also, the ABP, where selected hematological variables (Hb and RET%) are longitudinally monitored to create individually calculated thresholds can be applied to detect blood doping [50]. It has been suggested that miRNAs can function as complementary biomarker in ABP and miR486-5p, miR-486-3p and miR144-3p have all been suggested as biomarkers for blood doping [20-22]. Our results show no association between miR144 and any of the ABP hematological parameters, whereas a minor correlation between, miR486-5p and Hb was discerned, further enhancing a connection between these two markers. However, to function as a longitudinal passport marker, the expression profile should show low intra-individual variations. Surprisingly, the stability of these miRNAs has not been longitudinally evaluated before, and the relatively large intra-subject variations observed herein may be a concern. Moreover, administration studies of long acting erythropoietin stimulating agents (MIRCERA) both in humans [20] and in equines [51], reveal that miRNA-144 increase is not seen in all subjects implying that miRNA as a biomarker in anti-doping context may be less useful.

### 4.5 Limitations and strengths

One limitation in our study is the small sample size (n=13) and hence minor hormone-mediated effects on the miRNAs may not be found. Moreover, some values were missing, due to availability of samples and detection rate, and consequently three samples (one from each menstrual phase) per subject could not be included which decreases the power of our statistical analysis. For miRNAs in the circulation, either miR-5p or miR-3p is preferentially active [2]. Unfortunately, not all studies declare which miRNA strand is studied and thus comparing the results from our study, with other studies may be difficult.

Nevertheless, our study population is the largest population in terms of studying these miRNAs in relation to the menstrual cycle. Furthermore, a challenge in miRNA analyses is to identify the proper control miRNAs for normalization [52]. In this study, we show that the two housekeeping miRNAs used for normalization were not affected by the menstrual phases and that the use of only miR26b provided similar results as when miR26b and miR27 were combined.

### 4.6 Future studies

In the future, it is important to continue to study other putative miRNA diagnostic markers in relation to the menstrual cycle, to understand if the expression profile may fluctuate [31]. Not considering menstrual cycle phases when studying biomarkers in fertile women may introduce bias [53-56]. The association between miRNAs, EPO, hematopoiesis, lipids and atherosclerosis should be further explored, in both men and women. Moreover, to evaluate the utility of novel biomarkers, studies are ideally performed in healthy volunteers prior to investigations in patient populations. To further investigate the associations found in this study, a similar investigation should be performed in men.

## 5. Conclusion

To summarize, the results from this explorative pilot study, suggest that EPO, miR-144-3-p and miRNA-486-5p in blood, do not vary during the menstrual cycle, and thus may be used as biomarkers in women without considering the menstrual cycle phases. Nevertheless, both miR144 and miR486 showed large intra-individual variations and hence their use as longitudinal biomarkers might not be optimal. Moreover, miRNAs and EPO were associated to lipids and markers of hematopoiesis-miR144-3p was positively associated with HDL-C, whereas miR-486-5p was associated to EPO, red blood cells and hemoglobin. Serum erythropoietin showed significant correlations to lymphocytes, hemoglobin, high-density cholesterol and triglycerides. The association between miRNAs, EPO and other biomarkers of atherosclerosis should be further explored in the future.

## Abbreviations

(EPO): Erythropoietin
(rhEPO): recombinant human EPO
(ESA): erythropoietin stimulating agents
(RBC): red blood cell count
(RET): reticulocyte
(Athlete Biological Passport): ABP
(miRNA): microRNA
(Hb): hemoglobin
(HDL-C): high-density-cholesterol
(LDL-C): low-density-cholesterol
(TG): triglyceride
(CVD): cardiovascular disease
(AMI): acute myocardial infarction
(ABT): autologous blood transfusion

## Acknowledgements

The authors would like to express their sincere gratitude to all healthy volunteers who participated in the study. The authors also wish to express their gratitude to study nurse Susanne Broström at the Anti-Doping Hotline, Department of Pharmacology, Karolinska University Hospital for skillful work with the study and for logistical support.

## Ethics approval statement

The study was approved by the Human Ethics Committee of Karolinska Institutet (Dnr 2018/481-31/2). The study was performed in accordance with the Declaration of Helsinki. Written informed consent was obtained from all patients prior to inclusion in the study.

## Author contributions

Helena Bergström: data curation, statistical analysis, visualization, second draft of manuscript and onwards, rewriting and editing of manuscript. Lena Ekström: project administration, conceptualization, funding application, application to Human Ethics Committee of Karolinska Institutet, Biobank of Sweden and Swedish Data Protection Authority. Recruiting and screening of subjects. Laboratory analysis of miRNA. Statistical analyses and visualization. Writing first rough draft and reviewing manuscript. Supervision. Linda Björkhem-Bergman: conceptualization, investigation of subjects (study physician), writing-editing and reviewing. Carmel Heiland: method and analysis of EPO, writing-editing and reviewing. All authors contributed to the manuscript and read and approved the final manuscript.

## Data availability statement

As sample size is small, there is a risk for the individual privacy being compromised. De-identified data are therefore available from the corresponding author upon reasonable requests.

## References

1. Pritchard, C.C., H.H. Cheng, and M. Tewari, MicroRNA profiling: approaches and considerations. Nat Rev Genet, 2012. 13(5): p. 358–69.

2. Boon, R.A. and S. Dimmeler, MicroRNA-126 in atherosclerosis. Arterioscler Thromb Vasc Biol, 2014. 34(7): p. e15–6.

3. Valadi, H., et al., Exosome-mediated transfer of mRNAs and microRNAs is a novel mechanism of genetic exchange between cells. Nat Cell Biol, 2007. 9(6): p. 654–9.

4. Kim, K.M., et al., RNA in extracellular vesicles. Wiley Interdiscip Rev RNA, 2017. 8(4).

5. Michell, D.L. and K.C. Vickers, Lipoprotein carriers of microRNAs. Biochim Biophys Acta, 2016. 1861(12 Pt B): p. 2069–2074.

6. Wakabayashi, I., et al., Blood levels of microRNAs associated with ischemic heart disease differ between Austrians and Japanese: a pilot study. Sci Rep, 2020. 10(1): p. 13628.

7. de Lucia, C., et al., microRNA in Cardiovascular Aging and Age-Related Cardiovascular Diseases. Front Med (Lausanne), 2017. 4: p. 74.

8. Kuokkanen, S., et al., Genomic profiling of microRNAs and messenger RNAs reveals hormonal regulation in microRNA expression in human endometrium. Biol Reprod, 2010. 82(4): p. 791–801.

9. Florijn, B.W., et al., Gender and cardiovascular disease: are sex-biased microRNA networks a driving force behind heart failure with preserved ejection fraction in women? Cardiovasc Res, 2018. 114(2): p. 210–225.

10. Goretti, E., D.R. Wagner, and Y. Devaux, miRNAs as biomarkers of myocardial infarction: a step forward towards personalized medicine? Trends Mol Med, 2014. 20(12): p. 716–25.

11. Cipollone, F., et al., A unique microRNA signature associated with plaque instability in humans. Stroke, 2011. 42(9): p. 2556–63.

12. Sharma, B., et al., Expression of miR-18a-5p, miR-144-3p, and miR-663b in colorectal cancer and their association with cholesterol homeostasis. J Steroid Biochem Mol Biol, 2021. 208: p. 105822.

13. Vekic, J., et al., Obesity and dyslipidemia. Metabolism, 2019. 92: p. 71–81.

14. Ramirez, C.M., et al., Control of cholesterol metabolism and plasma high-density lipoprotein levels by microRNA-144. Circ Res, 2013. 112(12): p. 1592–601.

15. Chen, B., et al., Altered Plasma miR-144 as a Novel Biomarker for Coronary Artery Disease. Ann Clin Lab Sci, 2018. 48(4): p. 440–445.

16. Bye, A., et al., Circulating microRNAs predict future fatal myocardial infarction in healthy individuals - The HUNT study. J Mol Cell Cardiol, 2016. 97: p. 162–8.

17. Wang, L.S., et al., MicroRNA-486 regulates normal erythropoiesis and enhances growth and modulates drug response in CML progenitors. Blood, 2015. 125(8): p. 1302–13.

18. Zhang, R., et al., Expression of circulating miR-486 and miR-150 in patients with acute myocardial infarction. BMC Cardiovasc Disord, 2015. 15: p. 51.

19. Wakabayashi, I., Y. Sotoda, and R. Eguchi, Relationships among erythrocyte-derived microRNAs in serum of healthy donors. Clin Chim Acta, 2020. 507: p. 7–10.

20. Leuenberger, N., et al., Circulating microRNAs as long-term biomarkers for the detection of erythropoiesis-stimulating agent abuse. Drug Test Anal, 2011. 3(11-12): p. 771–6.

21. Gasparello, J., et al., Altered erythroid-related miRNA levels as a possible novel biomarker for detection of autologous blood transfusion misuse in sport. Transfusion, 2019. 59(8): p. 2709–2721.

22. Leuenberger, N., N. Robinson, and M. Saugy, Circulating miRNAs: a new generation of antidoping biomarkers. Anal Bioanal Chem, 2013. 405(30): p. 9617–23.

23. Libby, P., The changing landscape of atherosclerosis. Nature, 2021. 592(7855): p. 524–533.

24. Zhang, M., et al., Erythropoietin promotes abdominal aortic aneurysms in mice through angiogenesis and inflammatory infiltration. Sci Transl Med, 2021. 13(603).

25. Wu, B.W., et al., Downregulation of microRNA-135b promotes atherosclerotic plaque stabilization in atherosclerotic mice by upregulating erythropoietin receptor. IUBMB Life, 2020. 72(2): p. 198–213.

26. Bergstrom, H., et al., Variations in biomarkers of dyslipidemia and dysbiosis during the menstrual cycle: a pilot study in healthy volunteers. BMC Womens Health, 2021. 21(1): p. 166.

27. Mullen, J., et al., Fluctuations of hematological Athlete Biological Passport biomarkers in relation to the menstrual cycle. Drug Test Anal, 2020.

28. Schulze, J., et al., Urinary steroid profile in relation to the menstrual cycle. Drug Test Anal, 2021. 13(3): p. 550–557.

29. Livak, K.J. and T.D. Schmittgen, Analysis of relative gene expression data using real-time quantitative PCR and the 2(-Delta Delta C(T)) Method. Methods, 2001. 25(4): p. 402–8.

30. Friedewald, W.T., R.I. Levy, and D.S. Fredrickson, Estimation of the concentration of lowdensity lipoprotein cholesterol in plasma, without use of the preparative ultracentrifuge. Clin Chem, 1972. 18(6): p. 499–502.

31. Rekker, K., et al., Circulating microRNA Profile throughout the menstrual cycle. PLoS One, 2013. 8(11): p. e81166.

32. Ekström, L., et al., miRNA-27b levels are associated with CYP3A activity in vitro and in vivo. Pharmacol Res Perspect, 2015. 3(6): p. e00192.

33. Kim, M., et al., MIR144 and MIR451 regulate human erythropoiesis via RAB14. Br J Haematol, 2015. 168(4): p. 583–97.

34. Vu, L., et al., Analysis of Argonaute 2-microRNA complexes in ex vivo stored red blood cells. Transfusion, 2017. 57(12): p. 2995–3000.

35. Niculescu, L.S., et al., MiR-486 and miR-92a Identified in Circulating HDL Discriminate between Stable and Vulnerable Coronary Artery Disease Patients. PLoS One, 2015. 10(10): p. e0140958.

36. van Vliet, T., F. Casciaro, and M. Demaria, To breathe or not to breathe: Understanding how oxygen sensing contributes to age-related phenotypes. Ageing Res Rev, 2021. 67: p. 101267.

37. Makinoda, S., et al., Erythropoietin, granulocyte-colony stimulating factor, interleukin-1 beta and interleukin-6 during the normal menstrual cycle. Int J Gynaecol Obstet, 1996. 55(3): p. 265–71.

38. Alwachi, S.N. and A.J.H. Al-Saedi, Effect of Erythropoietin Therapy on the Lipid Profile in Patients with Chronic Kidney Disease–A Single Center Study. Journal of Nephrology & Therapeutics, 2013. 03(03).

39. Siamopoulos, K.C., et al., Long-term treatment with EPO increases serum levels of highdensity lipoprotein in patients with CKD. Am J Kidney Dis, 2006. 48(2): p. 242–9.

40. Li, J., et al., Kidney-secreted erythropoietin lowers lipidemia via activating JAK2-STAT5 signaling in adipose tissue. EBioMedicine, 2019. 50: p. 317–328.

41. Tóthová, Z., et al., STAT5 as a Key Protein of Erythropoietin Signalization. Int J Mol Sci, 2021. 22(13).

42. Christensen, B., et al., Erythropoietin does not activate erythropoietin receptor signaling or lipolytic pathways in human subcutaneous white adipose tissue in vivo. Lipids Health Dis, 2016. 15(1): p. 160.

43. Noguchi, C.T., Erythropoietin regulates metabolic response in mice via receptor expression in adipose tissue, brain, and bone. Exp Hematol, 2020. 92: p. 32–42.

44. Suresh, S., et al., Erythropoietin-Induced Changes in Bone and Bone Marrow in Mouse Models of Diet-Induced Obesity. Int J Mol Sci, 2020. 21(5).

45. Cao, Y., Erythropoietin in cancer: a dilemma in risk therapy. Trends Endocrinol Metab, 2013. 24(4): p. 190–9.

46. Cantarelli, C., A. Angeletti, and P. Cravedi, Erythropoietin, a multifaceted protein with innate and adaptive immune modulatory activity. Am J Transplant, 2019. 19(9): p. 2407–2414.

47. Ketelhuth, D.F. and G.K. Hansson, Adaptive Response of T and B Cells in Atherosclerosis. Circ Res, 2016. 118(4): p. 668–78.

48. Suresh, S., P.K. Rajvanshi, and C.T. Noguchi, The Many Facets of Erythropoietin Physiologic and Metabolic Response. Front Physiol, 2019. 10: p. 1534.

49. Martin, L., et al., Improved detection methods significantly increase the detection window for EPO microdoses. Drug Test Anal, 2021. 13(1): p. 101–112.

50. Malcovati, L., C. Pascutto, and M. Cazzola, Hematologic passport for athletes competing in endurance sports: a feasibility study. Haematologica, 2003. 88(5): p. 570–81.

51. Loup, B., et al., miRNAs detection in equine plasma by qPCR for doping control: assessment of blood sampling and study of eca-miR-144 as potential ESA biomarker. Drug Test Anal, 021.

52. McDonald, J.S., et al., Analysis of circulating microRNA: preanalytical and analytical challenges. Clin Chem, 2011. 57(6): p. 833–40.

53. Schisterman, E.F., S.L. Mumford, and L.A. Sjaarda, Failure to consider the menstrual cycle phase may cause misinterpretation of clinical and research findings of cardiometabolic biomarkers in premenopausal women. Epidemiol Rev, 2014. 36: p. 71–82.

54. Callegari, E.T., et al., Bone turnover marker reference intervals in young females. Ann Clin Biochem, 2017. 54(4): p. 438–447.

55. Lau, E.S., et al., Sex Differences in Circulating Biomarkers of Cardiovascular Disease. J Am Coll Cardiol, 2019. 74(12): p. 1543–1553.

56. Vashishta, S., S. Gahlot, and R. Goyal, Effect of Menstrual Cycle Phases on Plasma Lipid and Lipoprotein Levels in Regularly Menstruating Women. J Clin Diagn Res, 2017. 11(5): p. CC05–CC07.

